# CArP - CApture-based sequencing for Pathogen surveillance in complex matrices

**DOI:** 10.1101/2025.04.20.648581

**Authors:** Paul B. Stege, Frank Harders, Alex Bossers, Michael S. M. Brouwer

## Abstract

This study highlights the development and application of a novel capture-based long-read sequencing approach, CArP, designed to benefit pathogen surveillance. This combines targeted gene enrichment with the contextual insights of long-read sequencing, providing a highly efficient and flexible approach for pathogen detection. In this initial version, antimicrobial resistance (AMR) genes are utilized as marker genes that improve sequencing sensitivity. This allowed the detection of bacteria in broiler caecal content samples that carried resistance genes for critical antibiotics, including macrolides and quinolones. These specific AMR genes would have remained undetected when traditional metagenomic shotgun sequencing would have been applied, as it failed to detect these genes and therefore putative pathogens. This demonstrates the potential to detect and characterize pathogens more effectively compared to traditional sequencing strategies. The modular design of CArP furthermore demonstrates expansion with additional marker genes, such as viral or fungal markers, that can enable broader pathogen coverage. Taken together, this approach allows to advance pathogen detection and can contribute to global efforts in pandemic preparedness and the subsequent response.

## Introduction

Effective surveillance of pathogens is essential for safeguarding public health and preventing the emergence and spread of infectious diseases. Traditional culture-based sequencing methods are highly specific due to selective culturing, but costly when screening for large numbers of pathogens. Culture-free sequencing offers broader pathogen detection but suffers from a lack of specificity to detect pathogens among large quantities of other microbes. This can partially be compensated by increasing sequencing depth, at the risk of becoming costly again and pathogens still going undetected.

Given these limitations, there is a need to advance pathogen surveillance by a method that has a high specificity, but with the scalability and detection range of culture-free approaches. To address this, we developed a culture-free approach for pathogen surveillance called ‘Capture-based sequencing for Pathogen surveillance’ (CArP), which combines probe-based capturing techniques with long-read Oxford Nanopore Technology sequencing. This approach applies probes to target marker genes, allowing for the selective capture of DNA fragments for long-read culture-free sequencing. By analysing DNA regions flanking these marker genes, species origin can be determined and pathogens can be identified. The selection of marker genes will determine the degree of specificity over conventional culture-free approaches. In this first version, antimicrobial resistance (AMR) genes are used as marker genes. While AMR genes are not a marker of pathogens, pathogens often carry a wide range of AMR genes (Juhas e.a. 2009; Radovanovic e.a. 2023; Rice 2008; Rozwandowicz e.a. 2018). The marker gene panel allows modular expansion with additional gene markers, that target a wider range of bacteria or even fungal and viral pathogens. This flexibility is essential for addressing emerging infectious threats and enhancing global pandemic preparedness.

In this pilot study, we apply CArP to complex matrices and access its potential for pathogen surveillance. Its performance is compared to that of traditional metagenomic shotgun sequencing (MGS), demonstrating potential limitations of alternative screening strategies and highlighting the advantage of CArP. This approach therefore forms a basis for advancing pathogen surveillance, which is essential for pandemic preparedness and the subsequent response.

## Results

To evaluate the effectiveness of our CArP approach compared to traditional metagenomic shotgun sequencing (MGS), we used broiler cecum content samples as representative of complex microbial communities. Eight pooled caecal samples from individual farms, representing 10 broilers each, were used for this comparison. CArP utilizes antimicrobial resistance (AMR) genes to enrich target DNA sequences, we therefore first assessed differences in AMR gene detection between CArP and traditional MGS.

### Effectiveness of CArP in AMR gene detection

After read preprocessing, MGS resulted in a median of 158.5M reads per sample, of which 3.60E-2% corresponded to AMR genes (table 1). After read preprocessing, CArP resulted in 908 K reads per sample of which 36.4% corresponded to AMR genes. These reads had an N50 of 658 bases [SEM 3 bases]. The WHO classification system was used to further categorize antimicrobial resistance (AMR) genes regarding their importance, based on their association with pathogens and impact on human health (World Health Organization 2019). This resulted in the following two groups: (1) priority AMR genes and (2) lower-priority AMR genes (table s1).

**Table 1:**
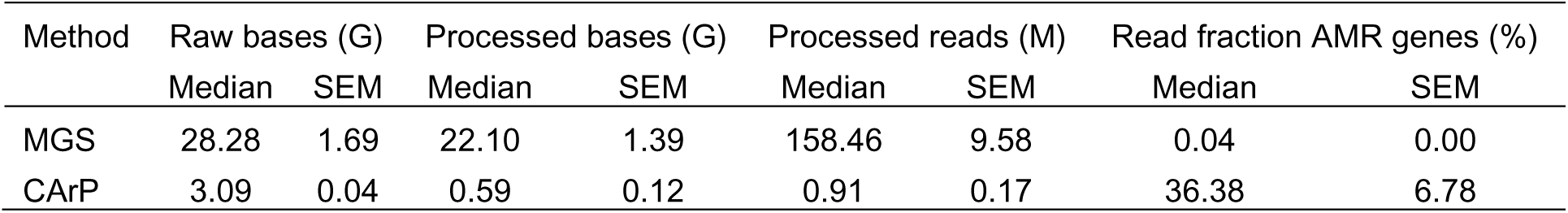
Comparison of Metagenomic Shotgun Sequencing and CArP sequence output and AMR-aligned read fraction.

| Method | Raw bases (G) |  | Processed bases (G) |  | Processed reads (M) |  | Read fraction AMR genes (%) |  |
| --- | --- | --- | --- | --- | --- | --- | --- | --- |
|  | Median | SEM | Median | SEM | Median | SEM | Median | SEM |
| MGS | 28.28 | 1.69 | 22.10 | 1.39 | 158.46 | 9.58 | 0.04 | 0.00 |
| CArP | 3.09 | 0.04 | 0.59 | 0.12 | 0.91 | 0.17 | 36.38 | 6.78 |

In both the MGS and CArP, the lower-priority AMR genes constituted the majority, with a median relative abundance of 77.9% [SEM 1.2%] and 74.0% [SEM 2.1%], respectively, of all detected AMR reads. For both methods, the most abundant resistance gene class in this group was tetracycline, with a median relative abundance of 72.1% [SEM 1.7%] for MGS and 68.8% [SEM 2.9%] for CArP. Most notably, CARP was able to detect two additional gene classes; macrolide resistance genes with a relative abundance of 0.04% [SEM 0.01%] and quinolone resistance genes with a relative abundance of 0.05% [SEM 0.02%]. We additionally performed capture-based sequencing while targeting only the critically important AMR genes to further increase sensitivity (figure 1c). We refer to this as the CArP-priority setup, as opposed to the CArP-complete setup that targets all AMR genes. The CArP-priority setup resulted in the increased proportion of sequenced priority AMR genes from 26.0% [SEM 2.1%] to 74.8 [SEM 1.9%], compared to the CArP-complete setup.

**Figure 1:**
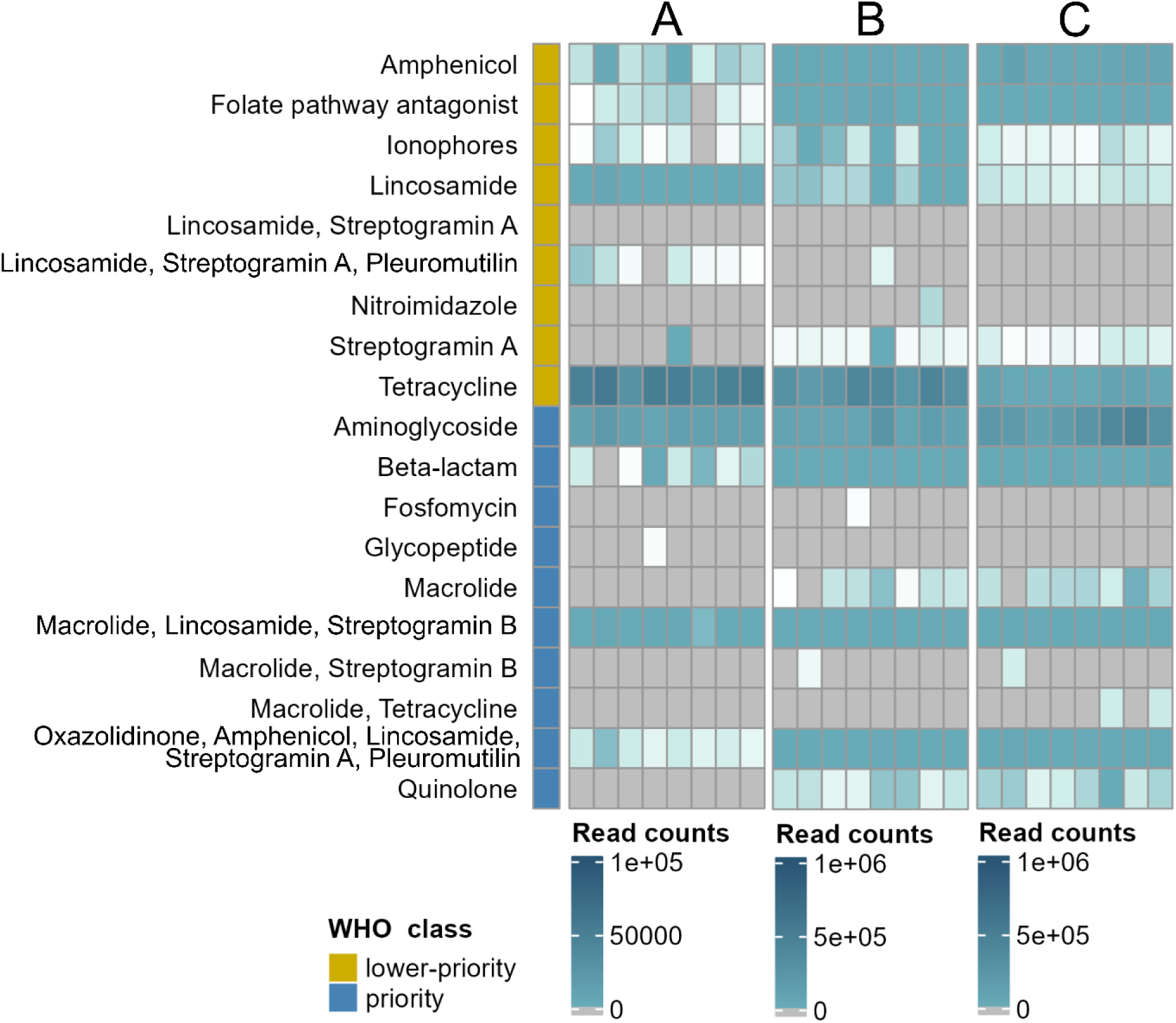
Abundance of AMR gene classes per sequencing method. Each row represents a sample, while panels represent the three applied sequencing methods: (A) metagenomic shotgun sequencing (MGS), (B) CArP using the complete probe set and (C) CArP with the priority probe set.

AMR gene detection was further analysed by visualizing AMR genes within each classification group as a function of sequencing depth. Due to high variability in gene complexity within the applied ResFinder AMR database, where some genes differ by a single nucleotide, a 90% sequence identity threshold was applied. In the downstream analysis, we report AMR genes, which technically refer to these gene representatives, allowing for efficient comparison (figure 2; table s2).

**Figure 2:**
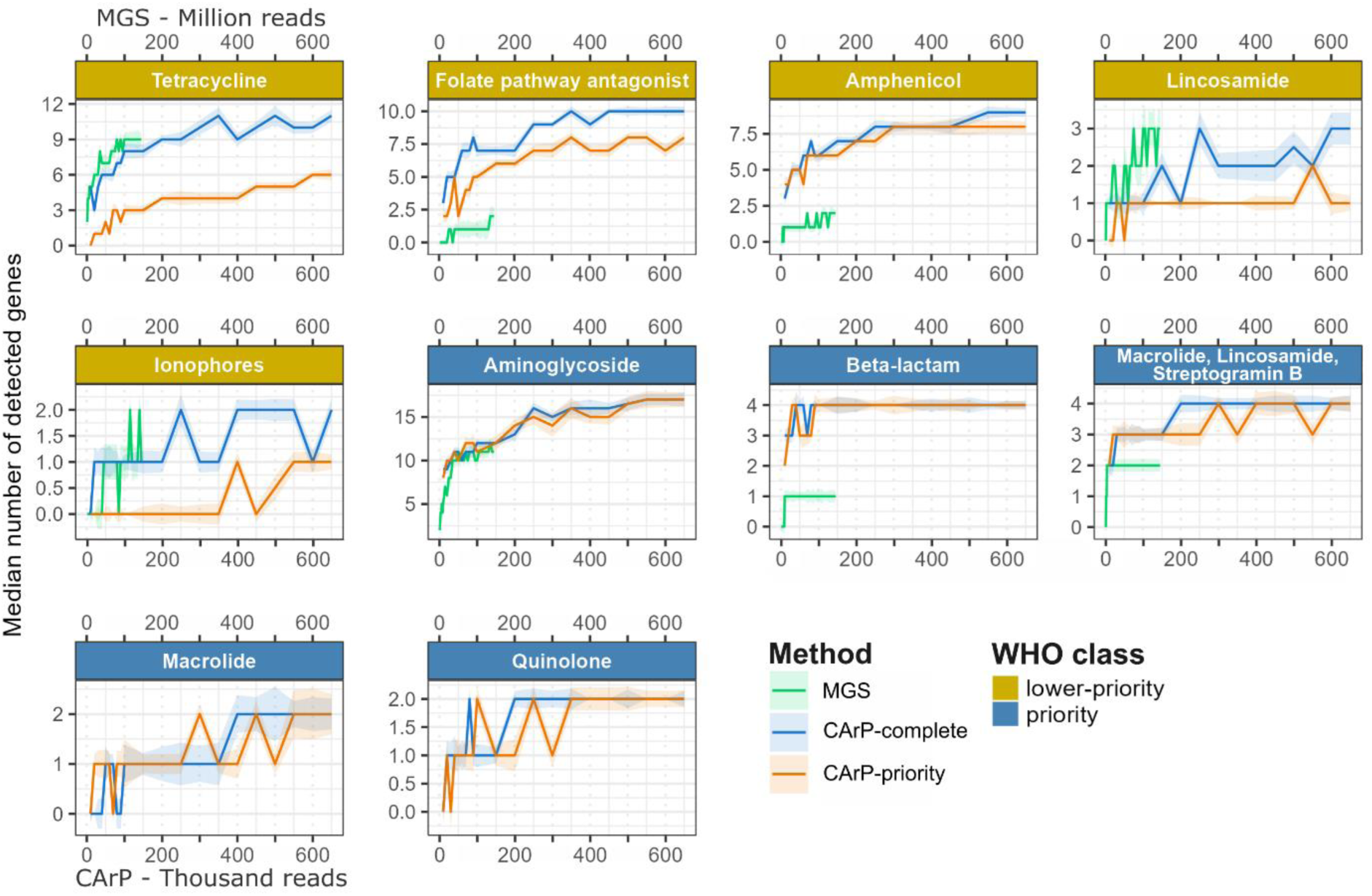
Number of detected antimicrobial resistance (AMR) genes per sequencing method. AMR genes are clustered into representative genes at 90% identity. The median number of AMR genes per sequencing method is represented by lines, with hue indicating the standard error of the mean (see legend). Panels represent AMR gene classes, where lower-priority AMR genes are coloured in gold, and priority AMR genes in blue. The x-axis is number of reads (thousands for ONT, and million for MGS).

At a relatively high sequencing depth of 145M reads per sample, MGS detected a high number of aminoglycoside resistance genes (median of 11 genes per sample) and tetracycline resistance genes (median of 9 genes per sample). At a sequence depth of 20M reads per sample, this rapidly reduced to a median of 6 aminoglycoside and 6 tetracycline resistance genes. While the difference in methodology between MGS and CArP prevent a fair comparison, we can conclude that using the CArP-complete setup detects more resistance genes for the classes: aminoglycoside; tetracycline; folate pathway antagonist; amphenicol; beta-lactam; macrolide, lincosamide, streptogramin B. Moreover, CArP is able to detect macrolide and quinolone resistance genes, whereas MGS does not. For the classes lincosamide ionophores the results fluctuate, but both CArP and MGS seem to detect an equal number of resistance genes at their maximum sequencing depth (3 lincosamide and 2 ionophore resistance genes). The CArP-priority probe set is not designed to detect all AMR genes, whereas the CArP-complete set is. We therefore expected the observed reduction in AMR gene detection for the classes: tetracycline, folate pathway antagonist, lincosamide and ionophores (figure 2). There is, however, no clear improvement in the detection of priority AMR genes at a lower sequencing depth (figure 2).

A major difference between MGS and the CArP approach, is the amount of information stored on long reads. We therefore used the regions flanking the detected AMR genes to predict species origin.

### AMR gene flanking region analysis for species and plasmid prediction

Reads containing macrolide or quinolone resistance genes were used to visualize gene context. DNA regions flanking AMR genes were selected from sequence reads and used to predict species origin and plasmid association (figure 3).

**Figure 3:**
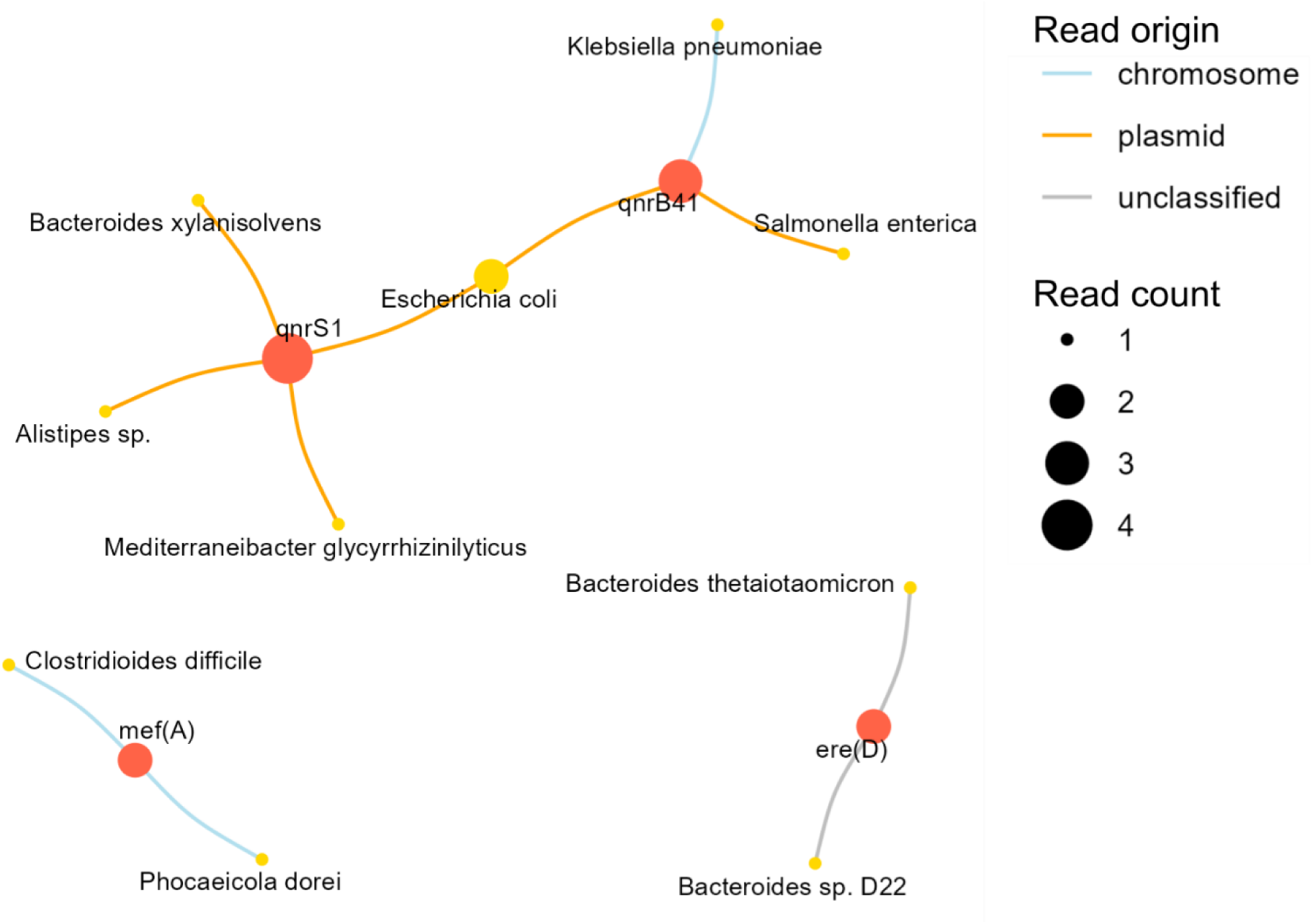
Sequence read-based relationship between AMR genes and prediction of species and plasmid origin. Nodes are coloured to indicate species (yellow) or AMR genes (red). Node size represents the number of reads matching with each hit. The connecting lines (edges) represent a species and AMR gene that are detected on a single read. Edges are coloured according to plasmid prediction.

The erythromycin resistance gene *ere(D)* contained flanking regions that matched *Bacteroides thetaiotaomicron* and *Bacteroides* sp. D22 (table s3). Regions flanking the macrolide resistance gene *mef(A)* matched *Clostridioides difficile* and *Phocaeicola dorei*. Regions flanking the quinolone resistance gene *qnrB41* matched *Salmonella enterica*-related plasmids, but also matched with non-plasmid entries of *Klebsiella pneumoniae* and *Escherichia coli* (figure 3, table s3). Regions flanking the second quinolone resistance gene, *qnrS1*, matched with *E*. *coli*-related plasmids, as well as non-plasmid entries of *Bacteroides xylanisolvens*, *Mediterraneibacter glycyrrhizinilyticus* and *Alistipes* sp.

To exemplify confirmation of read-based analyses, we additionally performed assembly-based analysis. For this, all *qnrB41 qnrS1* associated reads were assembled, resulting in 3 contigs: Contig 1 (1584 bp) contains *qnrS1* and flanking regions mainly match plasmid sequences from various Enterobacteriaceae species. Contig 2 (900 bp) includes *qnrB41* (table s3). It contains a small upstream flanking region that matches with *Helicobacter* and *Yersinia* species, while the larger downstream flanking region mainly matches *Escherichia coli* and *Salmonella enterica* derived plasmids. Contig 3 (927 bp) also carries *qnrB41*, with flanking regions matching plasmids from *E. coli* and species of *Klebsiella*, *Enterobacter* and other Enterobacteriaceae (table s3). From the MGS data we determined that the Enterobacteriaceae species detected though flanking region analysis have a low abundance of 0.096% [median relative abundance, SEM 0.034%], whereas all species belonging to Enterobacteriaceae make up 0.63% [median relative abundance, SEM 0.37%].

## Discussion

The findings of this study underscore the importance of advancing pathogen surveillance technologies and exemplifies limitations of alternative screening strategies such as metagenomic shotgun sequencing (MGS). We applied our proposed capture-based long-read sequence approach, CArP, to demonstrate the potential for efficient pathogen screening using marker genes. By combining the specificity of target enrichment with the gene-flanking insights provided by long-read sequencing, this approach enhances the effectiveness of pathogen screening. This first version applies antimicrobial resistance (AMR) genes as markers to increase specificity. We illustrate how this design offers flexibility via modular expansion, so that future versions can include additional genes targeting a wider range of pathogens. This approach can therefore form the basis of a more effective modular pathogen surveillance, essential for pandemic preparedness and response.

Broiler caecal content samples were used as exemplary complex environmental samples. To prioritize pathogens that carry critically important AMR genes for human medicine, we designed two separate probe sets for capture-based sequencing. The CArP-complete set targets all AMR genes, whereas the CArP-priority set targets a subset of AMR genes, that are defined as critically important (World Health Organization 2019). This served two purposes; Firstly, the pooling of probe sets exemplifies modular expansion of probe sets. This will allow future expansion with additional marker genes such as novel AMR genes, toxin genes, or fungal and viral gene markers. Secondly, this setup allows to focus on a subset of priority AMR genes with an additional level of enrichment, referred to as the CArP-priority set. Using MGS data, we determined that the fraction of AMR genes in complex environmental DNA samples can be as low as 0.036%. The majority of these genes consist of tetracycline resistance genes and only a subset consists of priority AMR genes. Among these genes are those with the highest priority for human medicine, which include quinolone, beta-lactam and macrolide resistance genes. Notably, the CArP approach was able to detect quinolone and macrolide resistance genes in the broiler caecal content samples, while MGS did not detect these. Both CArP as MGS did detect *erm(B)*, *erm(F)* and *erm(G)* resistance genes, which are part of the combined macrolide-lincosamide-streptogramin B class. Despite the relatively high sequencing depth, MGS is likely to miss quinolone and macrolide resistance genes in the analysed samples, as their relative abundance is lower than 0.1% relative abundance. This highlights how MGS-based pathogen surveillance might fail to detect pathogens that carry these priority AMR genes, and exemplifies how it could fail te detect other low abundant targets of interest.

We furthermore compared the sensitivity of the CArP-complete, targeting all AMR genes to that of the CArP-priority, which only targets critically important AMR genes. The CArP-priority set achieved a 2.8-fold enrichment of sequenced priority AMR genes, increasing their relative abundance from 26.0% to 74.8%. This however did not directly translate in the increased detection sensitivity of low abundant gene classes like quinolone and macrolide resistance at for instance a lower sequencing depth. The incorporation of the large class of aminoglycoside resistance genes in the CArP-priority set might overshadow the enrichment effect, as aminoglycoside genes are highly abundant. Further exclusion of lower priority aminoglycoside resistance genes could help to further increase the sensitivity of low abundant priority AMR genes and the pathogens that carry these. Moreover, although the probe design currently includes an equal number of probes per gene, it is also possible to prioritize and boost certain marker genes.

To account for the high variability in gene complexity within the applied ResFinder AMR database, where some genes differ by a single nucleotide, and account for the applied 5% error tolerance during long-read processing, we clustered the database entries based on a 90% sequence identity threshold (Florensa e.a. 2022)(table s2). In our analysis, AMR genes technically refer to these gene representatives. Using this setup, we pinpointed quinolone and macrolide resistance genes that the CArP approach was able to detect but MGS did not. This concerned the priority AMR gene clusters that confer macrolide resistance *ere(D)* and *mef(A)*, in addition to quinolone resistance *qnrB41* and *qnrS1*. Flanking regions of the *ere(D)* gene matched with two *Bacteroides* species, neither of which are considered pathogenic. The *mef(A)* gene cluster represents 4 closely related *mef(A)* genes and flanking regions match *Phocaeicola dorei* and the potential pathogenic *Clostridioides difficile* (Buddle en Fagan 2023). The *qnrB41* gene cluster represents 52 closely related *qnrB* genes and the *qnrS1* cluster represents 11 highly similar *qnrS* genes (table s2). Using read-based analysis on flanking regions, we detected similarity to plasmids from *Salmonella enterica* and *Escherichia coli*, as well as fragments from *Klebsiella pneumoniae* and *Escherichia coli*. A subset of *qnrS1* reads also showed similarity to species of *Bacteroides xylanisolvens*, *Mediterraneibacter glycyrrhizinilyticus* and *Alistipes* sp. Among these, *Alistipes* sp. is described as an opportunistic pathogen (Parker et al., 2020). While read-based analysis allows for high specificity, assembly-based analysis allows to find consensus among reads. Assembly of all *qnrB41* and *qnrS1*-associated reads resulted in three contigs from which the larger flanking regions (120-654 bp) mainly matched Enterobacteriaceae derived plasmids. Optimization of the current short sequence reads (N50 = 658 bases) will improve the length of the flanking regions and allows improved species prediction. Plasmid prediction can be further improved by the successor of the currently used tool, PlasmidEC, since this was trained for single-isolate sequencing (Paganini e.a. 2024).

In two cases MGS detected genes that were not detected by CArP. This included the detection of glycopeptide resistance gene *VanHDX_6* in a single sample with 26 aligned sequence reads. This gene entry represents a VanD-type resistance operon. Either CArP did not detect this gene, or this gene was not present in the DNA fraction used for CArP due to its low abundance. In addition, MGS was able detect the *lsaE_1* resistance gene in several samples, whereas CArP detected this gene in a single sample. *lsaE_1* is part of the combined AMR resistance group of: lincosamide, streptogramin A and pleuromutilin. It is possible that the designed CArP probes target a suboptimal region, resulting in a reduced detection sensitivity. In this case, these probes should be optimized.

While sequencing costs are project-dependent, they will impact the overall benefit of the methods, alongside the methods’ sensitivity for pathogen detection. In this study, a single set of DNA samples was used for both MGS and capture-based sequencing. The sequencing depth for MGS was adjusted to 140M reads per sample, so that the costs of library prep, personnel and sequencing approximately matched the sequencing costs of CArP using a single probe set. This does not yet include the requires synthesis of probes. While probe synthesis will double the costs per sample for this experiment with eight sample pools, these probes would be sufficient for at least a thousand sample pools, greatly reducing the per-sample cost. Currently, 90% of capture-based sequenced reads are excluded during read preprocessing, due to short reads and low-quality reads. Optimization of read quality and therefore length will reduce sequencing costs and increase detection sensitivity. The issues with read quality are suspected to take place during processing steps downstream of the probe-hybridization step. The relatively short reads result in short regions flanking AMR genes, thus limiting the accuracy of species prediction. Therefore, the current protocol requires validation of detected pathogens, until the sequence read length is optimized. Despite these optimization points, the current version of CArP still outperforms MGS in terms of AMR-based pathogen detection. CArP could even be combined with qPCR of signature AMR genes such as *sul1* and *erm(B)* to enable quantitative detection, similar to the AMR detection used by Macedo et al. (Macedo e.a. 2021).

To conclude, we demonstrated how the developed approach can outperform MGS-based screening using gene marker-based pathogen detection. Final optimization points offer further potential to enhance its efficiency and sensitivity, advancing pathogen surveillance.

## Methods

### Sample collection, storage and DNA extraction

Broiler cecum content samples were obtained between January and March 2024 from slaughterhouses as part of the ongoing national monitoring for AMR as reported in Nethmap-MARAN (‘Nethmap MARAN’ 2024). Caecal content was collected from 10 individual broilers from a single slaughter batch, equal amounts per animal were pooled and mixed. Samples were snap-frozen in liquid nitrogen and transferred to storage at −80 °C. DNA was extracted from 0.2 g of caecal content using PureLink Microbiome DNA Purification Kit (A29789, Thermo Fisher Scientific, Waltham, MA, USA) according to the manufacturer’s instructions. Total DNA was quantified by using a ClarioStar (BMG Labtech, Ortenberg, Germany).

### Metagenomic shotgun sequencing and data processing

DNA samples were sent to GenomeScan B.V. (Leiden, the Netherlands) for Metagenomic shotgun sequencing. Library preparation was performed using the NEBNext® Ultra II DNA Library Prep kit (E7645S, NEB, Ipswich, USA) according to manufacturer’s protocols. Libraries were sequenced on a NovaSeq 6000 sequencer (Illumina, San Diego, USA) using S2 flow cells and the 2×150 bp paired-end kit (Illumina, San Diego, USA) according to company protocols. Sequencing reads were adapter-clipped, erroneous-tile filtered, and quality-trimmed at ≥Q20 (Phred score) using BBduk v38.96 and subsequently filtered for host DNA using the global-alignment algorithm of BBmap v38.96 with default settings and fast=t (ENSEMBLE broiler genome version esm107)(Bushnell 2013). Read pairs were then used for taxonomic classification by Kraken v2.1.2 using the premade standard Kraken RefSeq nucleotide database and applying a confidence cut-off of 0.1 (database version 12/01/2024)(Wood, Lu, en Langmead 2019). Antimicrobial resistance (AMR) gene analysis was conducted using BBmap v38.96 with global alignment, fast=t, and default settings along with a custom database (Bushnell 2013). This analysis uses the ResFinder database v2.2.1 (accessed on 2024-01-24) clustering at 90% identity using CD-hit v4.8.1 and settings *cd-hit-est -c 0.90 -n 8 -s 0.80* (table s2)(Fu e.a. 2012). Sequencing resulted in a median of 28.3 Gb [SEM 1.70 Gb] per sample (table 1). After host filtering and preprocessing, this was reduced to 22.1 Gb [SEM 1.39 Gb] per sample and 158.5M [SEM 9.6M] reads per sample. A median of 21.9% [SEM 7.3%] of these reads could be annotated to bacterial taxa and 3.6E-2% [SEM 3.7E-3%] to AMR genes.

### Target capture probe design

Antimicrobial resistance (AMR) genes were used as molecular markers, to selectively sequence longs strands of bacterial DNA and subsequentially determine if the detected species is a predicted pathogen. AMR genes were downloaded using the ResFinder database v2.2.1, accessed on 2024-01-24 (Florensa e.a. 2022). This database was split into: (1) priority AMR genes and (2) lower-priority AMR genes; according to the WHO List of critically important antimicrobials for human medicine (World Health Organization 2019)(table s1). For each set, the following steps were applied: (1) Sequence redundancy was reduced using the Dedupe tool from BBTools *minidentity=95*, generating a fasta file per gene sequence (Bushnell 2013). (2) Probes were then designed on each file, using Probetools v0.1.9 with settings *makeprobes -b 1 -T 90 -i 95 -c 100 -l 110 -c 90 -d 0 -m 1* (Kuchinski e.a. 2022). (3) The resulting probes were consequently used for redundancy analysis using Dedupe software with *settings minidentity=95*. (4) Probes were validated against the original ResFinder database using BLAT v35 with a 10% mismatch threshold, corresponding to a cutoff of at least 108 matching bases in a single block out of the 120-base probe length (Kent 2002) (table s4). This resulted in the final probe list, where a median of 1 probe binds each target, with a maximum of 4 probes per target (table s4). (5) Probe sets were sent to Agilent Technologies (Santa Clara, CA, USA) for synthesis (table s4). A fraction of both probe sets was pooled to create a set that captures all AMR genes, called the CArP-complete set. This was compared to the CArP-priority set, which only contains critically important AMR genes.

### Long-read capture-based sequencing and data alignment

DNA fragmentation was performed using the Covaris ME220 Focused-ultrasonicator (Covaris, Woburn, MA, USA). Fragmentation size was confirmed using the TapeStation (Agilent Technologies, Santa Clara, CA, USA). DNA probe-based enrichment was performed using the Agilent SureSelect XTHS 2DNA Reagent Kit (Agilent Technologies, Santa Clara, CA, USA), followed by amplification with KOD FX Neo (TOYOBO, Osaka, Japan; #KFX201). Post-amplification, DNA was purified using Dynabeads (Thermo Fisher Scientific, Waltham, MA, USA; # DB65305). Library preparation was performed using the native barcoding kit according to manufacturer’s protocols (Oxford Nanopore Technologies, Oxford, UK; #SQK-NBD114-96). Subsequently, libraries were sequenced on a Promethion P2S-01015 using R10.4.1 flow cells, according to company protocols. Base calling was performed using dorado, v0.6.2 using settings *demux --emit-fastq --kit-name* (Oxford Nanopore Technologies 2022). Sequencing reads were quality-trimmed at ≥Q13 (Phred score) and minimum length of 500bp using BBduk v39.01. AMR gene analysis was performed by first indexing the clustered ResFinder database (90% identity threshold) using Minimap2 v2.28-r1209 and *settings -x map-ont -k19 -w19 -t 24 -I100G* (Florensa e.a. 2022; Li 2018). This database was then used to identify AMR genes using Minimap2 and settings *-x map-ont -k19 -w19 -U50,500 -g10k, -m100*. Filtering in R was applied as follows: the harmonic mean selected the best gene hit for overlapping regions on a read, and genes with less than 60% coverage in a sample were excluded (Marić e.a. 2024).

Before taxonomical classification, processed reads were first masked for the regions that contained AMR genes. For this, Minimap2 was used to detect AMR genes with settings *-ax map-ont -k19 -w 19 -U50,500 -g10k -m 100 −2 -cs=short --sam-hit-only*, followed by conversion to a bed file using sam2bed v2.4.41 with settings *--do-not-sort -a* (Quinlan en Hall 2010). This alignment was then used to mask AMR regions on sequencing reads using bedtools v2.31.1 with settings *maskfasta* (Quinlan en Hall 2010). Consequently, masked reads were used for taxonomic classification using Minimap2 v2.28-r1209 and settings *-x map-ont -k19 -w 19 -U50,500 -g10k*, while using the NCBI Reference Sequence Database (accessed 09-10-2024). This contained 329,490 bacterial genomes that were downloaded using the NCBI datasets tool v16.23.0 (O’Leary e.a. 2024). After Minimap2 alignment, the harmonic mean was applied to select the top hit for each query sequence (Marić e.a. 2024). In parallel, PlasmidEC was performed on processed reads to predict chromosome or plasmid origin (Paganini e.a. 2024) using settings *-s general -f -l 500* and only the Centrifuge database output. Sequencing resulted in a median of 3092 Mb [SEM 36.8 Mb] per sample (table 1). After applying the adapter and quality filtering steps, this was reduced to 585 Mb [SEM 115 Mb] per sample and 908 K reads [SEM 170 K reads] per sample. These filtered reads had an N50 of 658 b [SEM 3 b] and 36.4% [SEM 6.78%] of the reads contained AMR genes and were therefore considered on-target. For assembly, reads harbouring *qnrB41* and *qnrS1* AMR genes were first extracted and used for assembly with Canu v2.2 and settings *-nanopore-raw genomeSize=20k minReadLength=400 minOverlapLength=250* (Koren e.a. 2017). Contigs were then used for alignment against the two quinolone resistance genes using Minimap2 and settings *-x map-ont -k19 -w 19 -U50,500 -g10k -m 100 −2*. The regions flanking these genes were then extracted and used for taxonomic annotation using BLAST and the bacterial core nucleotide database (Altschul e.a. 1990). The top 10 hits are reported (table s3).

### Data visualization

Visualization of sequencing data was performed in R v4.3.1, Rstudio v2023.09.1+494 (set seed 8591) and functions of R packages phyloseq v1.44. The relative abundance of detected AMR gene classes was calculated by applying taxonomic agglomeration on class level (tax_glom, phyloseq package) and plotted using functions of the microbiome v1.22 and ComplexHeatmap v2.16 packages (Lahti en Shetty 2017; Gu 2022). A cut-off of 5 reads was used to differentiate between AMR gene presence and absence. Rarefaction of ONT data was performed in steps of 1,000 reads for 10,000– 100,000 sub-samples and in steps of 50,000 reads for 150,000–650,000 sub-samples. For MGX data, rarefaction was conducted in steps of 1 million reads for 1–10 million sub-samples and in steps of 5 million reads for 15–145 million sub-samples. The rarefaction process applied the reformat function in BBTools with the *samplereadstarget* setting (Bushnell 2013). Sample 7 was excluded from this analysis due to a low read count in the ONT dataset (450,000 reads). Gene classes were subsequently visualized using ggplot2 v3.5.1 when at least two AMR genes were detected, based on the median number of detected (Wickham 2016). Sequence read-based relationship between AMR genes and prediction of species and plasmid origin was determined using exact matching of read IDs. Visualization uses packages igraph v1.2.6 (Csardi en Nepusz 2006).

## Supporting information

table_s1-Resfinder_WHO_classification

table_s2-ClusteredResFinderDB_90percent

table_s3-Quinolone_contig_blast

table_s4-Probe_design_BLAT_analysis

## Data availability

Raw sequencing data from metagenomic shotgun and Oxford Nanopore Technologies sequencing have been deposited in the NCBI Sequence Read Archive (SRA) under accession number PRJNA1188361. The phyloseq objects are available at: 10.5281/zenodo.14216441.

